# Population-level antagonism between FGF and BMP signaling steers mesoderm differentiation in embryonic stem cells

**DOI:** 10.1101/2023.03.24.534121

**Authors:** Marina Gattiglio, Michelle Protzek, Christian Schröter

## Abstract

The mesodermal precursor populations for different internal organ systems are specified during gastrulation by the combined activity of extracellular signaling systems such as BMP, Wnt, Nodal, and FGF. The BMP, Wnt and Nodal signaling requirements for the differentiation of specific mesoderm subtypes in mammals have been mapped in detail, but how FGF shapes mesodermal cell type diversity is not precisely known. It is also not clear how FGF signaling integrates with the activity of other signaling systems involved in mesoderm differentiation. Here, we address these questions by analyzing the effects of targeted signaling manipulations in differentiating stem cell populations with single cell resolution. We identify opposing functions of BMP and FGF, and map FGF-dependent and -independent mesodermal lineages. Stimulation with exogenous FGF boosts the expression of endogenous Fgfs while repressing Bmp ligands. This positive autoregulation of FGF signaling, coupled to the repression of BMP signaling, may contribute to the specification of reproducible and coherent cohorts of cells with the same identity via a community effect, both in the embryo and in synthetic embryo-like systems.

## Introduction

During the development of multicellular organisms, precursor cells for specialized organs need to differentiate in reproducible proportions and in spatially coherent domains. Although the main cell-cell communication systems that drive and coordinate differentiation in cell populations have been identified, how these systems interact in signaling networks is still not well understood.

The mesoderm encompasses precursor cells for important organ systems such as the blood, vasculature, heart, kidney, limbs, and the musculoskeletal system. In the mouse, the mesodermal precursor cells for these organ systems are specified during gastrulation from the pluripotent epiblast, a cup-shaped epithelium that is surrounded by extraembryonic tissues such as the extraembryonic ectoderm (ExE) and the visceral endoderm (Fig. 1A). Precursors of different mesoderm subtypes are found in distinct regions of the pre-gastrulation epiblast (Tam and Behringer, 1997), and traverse the primitive streak, the site of gastrulation, at different times. Cells from the proximal epiblast close to the ExE for example gastrulate first and will preferentially give rise to extraembryonic mesoderm and blood, followed by precursors of the heart and head mesoderm. In contrast, the paraxial and axial mesoderm, which encompasses precursor cells of the musculoskeletal system, arises later in more distal parts of the embryo (Fig. 1A, Ferretti and Hadjantonakis, 2019). Cell differentiation in the epiblast is thought to be governed by a system of signaling gradients, mainly generated by the extraembryonic cells that surround the epiblast (Arnold and Robertson, 2009; Tam and Loebel, 2007). The ExE for example expresses BMP4 ligands that establish a proximal-to-distal phosphorylation gradient of the BMP-signal transducer Smad1/5 (Arnold and Robertson, 2009; Morgani et al., 2018a; Winnier et al., 1995; Zhang et al., 2019). BMP signaling promotes the differentiation of proximal, but not distal mesoderm subtypes from pluripotent stem cells (Manfrin et al., 2019; Morgani et al., 2018a), consistent with a proposed role of the BMP signaling gradient for mesoderm patterning in the embryo (Tam and Loebel, 2007). The BMP signaling gradient is complemented by gradients of Wnt and Nodal signaling that promote the differentiation of more distal mesoderm and endoderm, respectively. The Wnt and Nodal gradients are established through a combination of localized ligand expression on the posterior side of the embryo, together with the secretion of signaling antagonists from the anterior visceral endoderm (AVE) at the embryo’s anterior side (Arnold and Robertson, 2009; Stower and Srinivas, 2018).

**Fig. 1:**
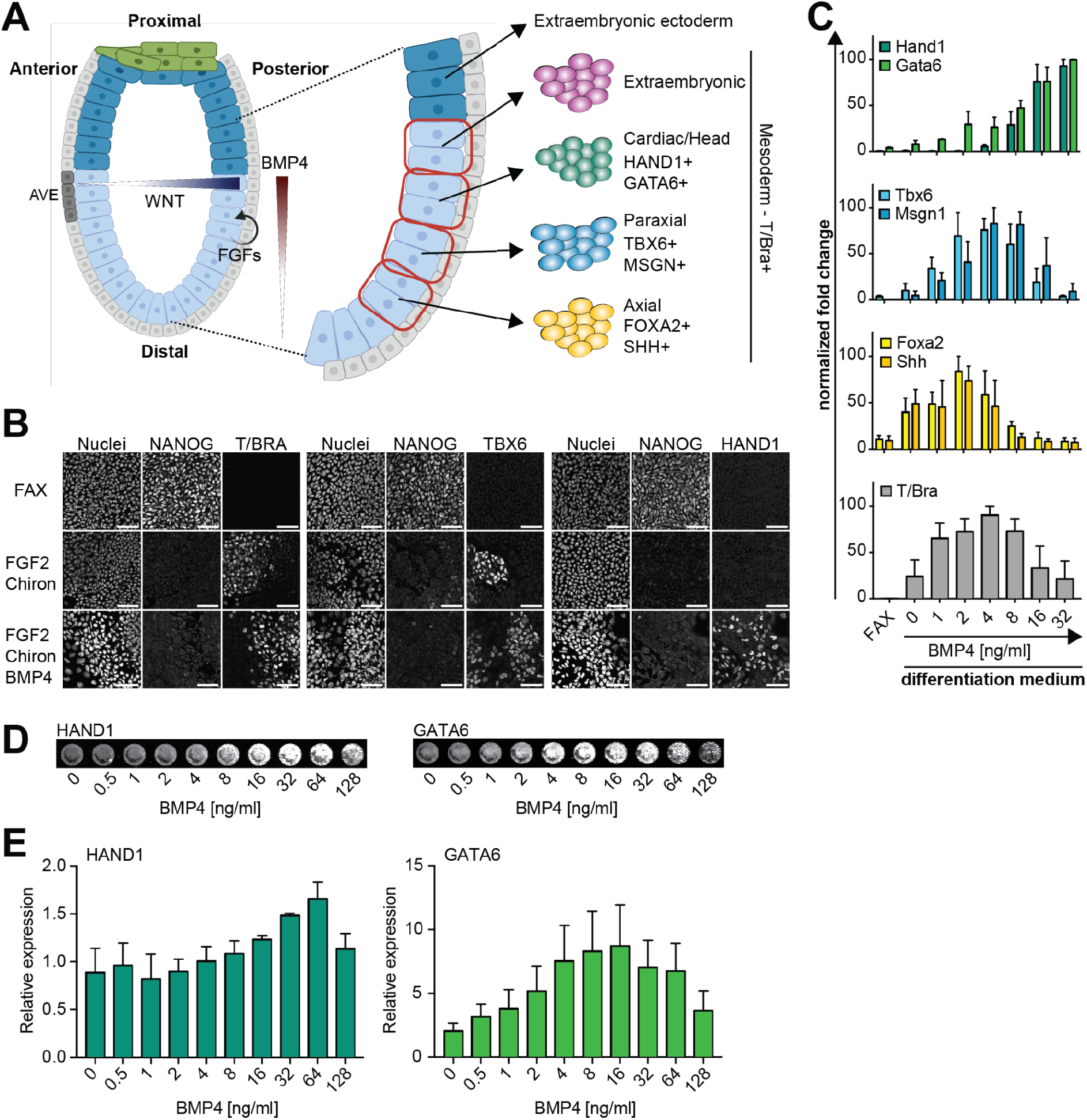
Concentration-dependent functions of BMP4 during mesoderm differentiation in vitro. **A** Schematic of signaling systems in the mouse embryo before gastrulation (left), and fate map of epiblast cells in the pre-gastrulation embryo (right). Marker genes for mesodermal subtypes are indicated. AVE, Anterior Visceral Endoderm. Created with BioRender.com. **B** Immunostaining for NANOG, T/BRA, TBX6, HAND1 of EpiSCs in FAX medium (top), or differentiated with FGF2 and Chi in the absence (middle) or presence (bottom) of BMP4. Nuclei were stained with Hoechst33342. Scale bars, 50 μm. **C** qPCR analysis of mesoderm markers in EpiSCs differentiated with a BMP4 concentration series. Measurements were normalized in each individual experiment by setting the highest expression value for each marker to 100. Plots show mean ± SEM from n=3 independent experiments. **D** Representative images from In-Cell Western detection of HAND1 and GATA6 in cells differentiated as in **C**. **E** Quantification of marker expression in the In-Cell Western experiment. Data for each marker and experiment was normalized to a negative control cultured without FGF and BMP. Error bars indicate SEM from n=3 independent experiments.

In addition to these three graded signaling systems, cell differentiation and patterning in the mesoderm critically relies on fibroblast growth factor (FGF) signaling. FGF signaling is most active in the primitive streak and the nascent mesoderm (Morgani et al., 2018b), mirroring the expression of Fgf8, Fgf4, Fgf3 and Fgf17 ligands in this region (Crossley and Martin, 1995; Maruoka et al., 1998; Niswander and Martin, 1992). Loss of Fgf signaling in Fgfr1- and Fgf8-mutant embryos impairs cell migration and leads to the accumulation of cells in the primitive streak (Ciruna and Rossant, 2001; Deng et al., 1994; Sun et al., 1999; Yamaguchi et al., 1994). Furthermore, although some proximal mesoderm forms in these mutants, the differentiation of more distal cell types such as the paraxial mesoderm is defective (Ciruna and Rossant, 2001; Deng et al., 1994; Sun et al., 1999; Yamaguchi et al., 1994). In human embryonic stem cells in contrast, FGF signaling has been proposed to be generally required for efficient mesoderm differentiation in cooperation with BMP signaling, since BMP4-treatment in the absence of FGF leads to the differentiation of extraembryonic cell types (Bernardo et al., 2011; Yu et al., 2011). It is thus an open question, exactly which mesodermal cell types are FGF-dependent, how FGF signaling is integrated with other signals present during mesoderm differentiation, and how this integration contributes to mesoderm patterning.

In recent years, in vitro models based on pluripotent cell populations that represent the epiblast have emerged as powerful tools to investigate the signaling control of cell differentiation and patterning during gastrulation. These systems allow testing the influence of individual factors in the absence of the complex environment of the embryo and its extraembryonic cell types (Morgani and Hadjantonakis, 2020). Even upon homogeneous external signaling cues, both 2D micropattern and 3D aggregate systems generate and spatially arrange diverse cell types that are normally found in the gastrulating embryo (Baillie-Benson et al., 2020; Brink et al., 2020; Morgani et al., 2018a). These observations suggest that, in addition to the signaling landscapes imposed by extraembryonic tissues, there exist epiblast-intrinsic mechanisms that orchestrate mesoderm differentiation and patterning.

Here we use a simple 2D epiblast stem cell (EpiSC) differentiation protocol to map concentration-dependent functions of BMP and FGF signaling during mesoderm differentiation. Through single cell sequencing and integration with published transcriptome datasets from the embryo, we show that FGF dose affects differentiation speed, and that it sets the proportions of discrete cell types in heterogeneous populations. The dose-dependent expression patterns of signaling genes suggest that FGF signaling is embedded in a positive autoregulatory loop, coupled to the repression of BMP signaling. This regulatory logic could establish an FGF-based community effect during mammalian mesoderm differentiation that contributes to generating spatially coherent groups of distal mesoderm cells that are spatially segregated from BMP-dependent proximal cell types.

## Results

### Opposing functions of BMP and FGF signaling during EpiSC differentiation

To obtain a homogeneous starting population for mesoderm differentiation, we cultured EpiSCs in medium containing ActivinA, FGF2, and the Wnt signaling inhibitor XAV939, which suppresses Wnt-induced cellular heterogeneity (FAX medium; Sumi et al., 2013; Tsakiridis et al., 2014). Under these culture conditions, cells were stained homogeneously positive for NANOG, but were negative for the pan-mesodermal marker T/BRA (Fig. 1B). We triggered general mesoderm differentiation following previously published protocols by exchanging ActivinA and XAV939 for 1 µM of the Wnt agonist Chir99021 (Chi) (Chal et al., 2015; Loh et al., 2014; Sudheer et al., 2016), and varied the concentration of BMP4 in this setting (Vallier et al., 2009). After three days of differentiation, both T/BRA and TBX6, a marker for distal mesodermal cell types, were expressed in the absence and presence of 10 ng/ml BMP4. In contrast, expression of HAND1, a marker for more proximal mesoderm, was observed only in BMP4-treated cultures (Fig. 1B). To determine more precisely how BMP4 concentration affects cell differentiation, we used qPCR of a panel of marker genes that show regionalized expression in the gastrulating embryo (Peng, 2016). Expression of Hand1 and Gata6, which are expressed in the posterior proximal part of the embryo, was highest at 16 and 32 ng/ml BMP4 (Fig. 1C, top). The expression of Tbx6 and Msgn1, which are expressed more distally in the embryo, peaked around 4 ng/ml BMP4, while the most distal markers Foxa2 and Shh peaked at even lower BMP4 concentrations (Fig. 1C). Expression of the pan-mesodermal marker T/Bra paralleled the expression of Tbx6 and peaked at 4 ng/ml BMP4. To corroborate these findings at the protein level, we performed an In-Cell Western assay where we further increased the concentration range of BMP4 (Fig. 1D, E). Even though the high cell densities required in an In-Cell Western could potentially impair the activity of BMP4 (Etoc et al., 2016), this experiment broadly confirmed the qPCR results. HAND1 expression increased for BMP4 concentrations higher than 8 ng/ml and peaked at 64 ng/ml BMP, whereas GATA6 expression increased across the whole BMP4 concentration range up to 16 ng/ml and then declined (Fig. 1D, E). Taken together, these results show that BMP promotes the expression of more proximal mesoderm markers, Hand1 and Gata6, while inhibiting markers of more distal fates, in a concentration-dependent manner. This is consistent with the proposed role of graded BMP signaling for cell type specification in the embryo.

Next, we set out to determine concentration-dependent effects of FGF signaling in the mesoderm differentiation protocol. We tested titration series of FGF2 and FGF4 in the presence of 8 ng/ml BMP, because previous experiments indicated that this BMP concentration gave a mixture of cell types. We first focused on T/Bra expression, using the T/Bra:mCherry signal in the dual Sox1/Brachyury SBR reporter cell line as a read-out (Deluz et al., 2016). For both FGF2 and FGF4, reporter expression levels increased up to a concentration of 12 ng/ml, the standard FGF2 concentration in EpiSC maintenance medium (Brons et al., 2007). While for FGF4, reporter expression further increased to a maximum of 77.9 ± 4.6% T/Bra-mCherry+ cells (mean ± SEM, n = 3 independent experiments) at a concentration of 96 ng/ml FGF4, for FGF2, the proportion of reporter-positive cells decreased for concentrations above 12 ng/ml (Fig. 2A, B). This biphasic effect of FGF2 is consistent with previous reports showing that high concentrations of FGF2 lead to reduced FGF/ERK signal transduction (Gharibi et al., 2020; Kanodia et al., 2014). However, because FGF2 is commonly used in published mesoderm differentiation protocols, and because the FGF2 and FGF4 showed similar behavior at concentrations up to 12 ng/ml, we focused on FGF2 for subsequent experiments. To determine how different FGF2 concentrations in this low range affect the expression of a range of mesodermal markers, we performed quantitative immunofluorescence. Consistent with results from the reporter cell line, the proportion of T/BRA-positive cells increased with FGF2 concentration, while the downregulation NANOG-expression occurred independently from FGF2 (Fig. 2C). The proportion of HAND1-positive cells was highest in the absence of FGF2, and decreased with increasing FGF2-dose (Fig. 2D), in contrast to TBX6 expression, which was detected only at FGF2-concentrations above 1.5 ng/ml (Fig. 2E). CDX2, a gene that is associated both with proximal, extraembryonic mesoderm, as well as with the paraxial mesoderm in the embryo (Beck et al., 1995; Morgani et al., 2018a), was expressed in the absence of FGF2, and its expression slightly increased at higher concentrations (Fig. 2D). The observation that FGF promotes the expression of the more distal mesoderm marker TBX6, and inhibits the expression of the more proximal marker HAND1 in a concentration-dependent manner suggests that FGF signaling opposes the function of BMP in the mesoderm differentiation protocol used here.

**Fig. 2:**
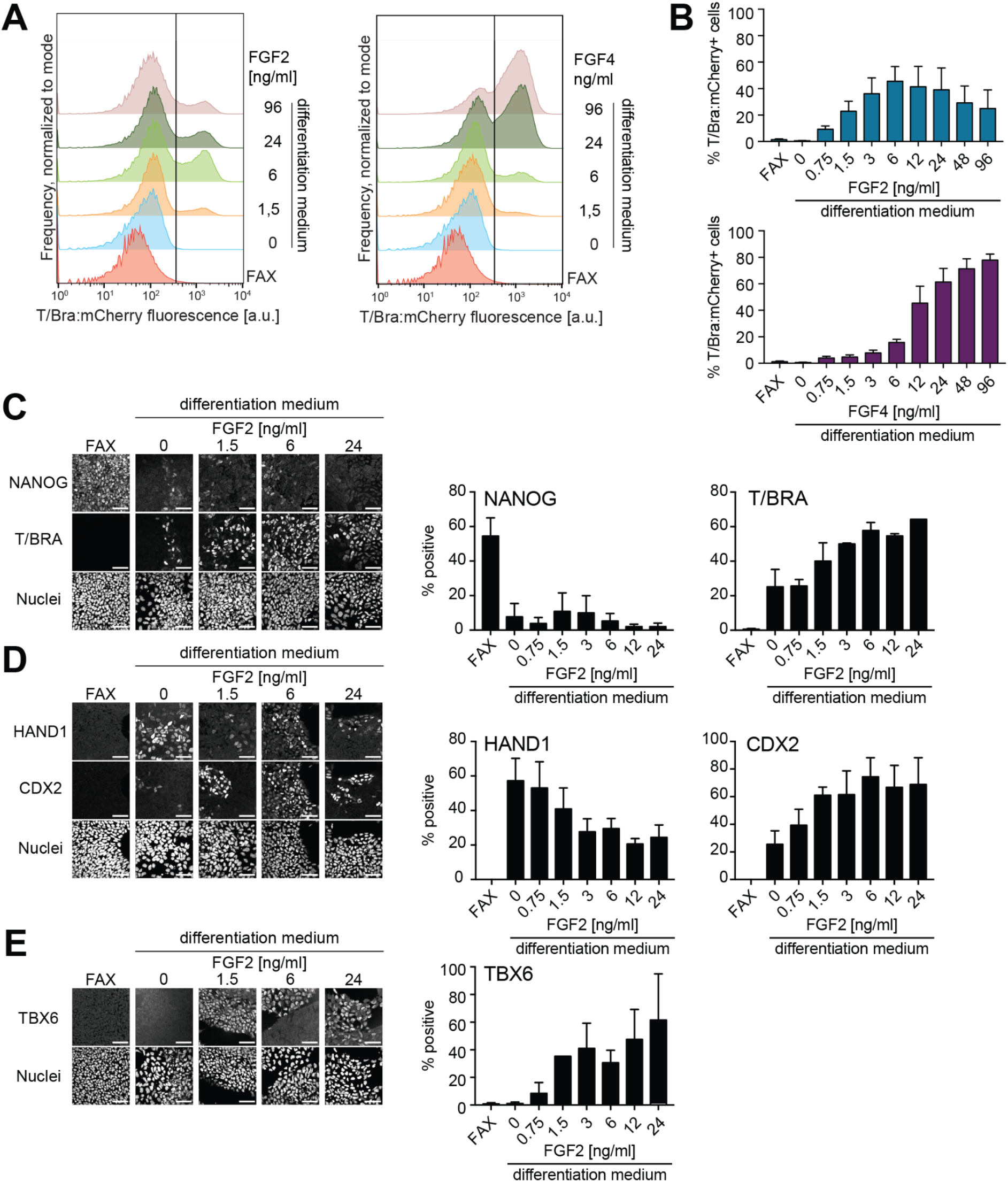
Concentration-dependent functions of FGFs during mesoderm differentiation in vitro. **A** Representative flow cytometry measurements of T/Bra: mCherry reporter expression in the SBR line differentiated with a range of FGF2 (left) or FGF4 concentrations (right). Vertical line indicates threshold between positive and negative cells. **B** Fraction of T/Bra:mCherry-positive cells measured by flow cytometry as in **A**. Data from N = 3 independent experiments, error bars indicate SEM. **C - E** Immunostaining for NANOG, T/BRA, HAND1, CDX2 and TBX6 (left) and quantification of the fraction of marker-positive cells (right) upon differentiation of EpiSCs with the indicated FGF concentration series. Cells were considered marker-positive when they belonged with >70% probability to the high-intensity component of a 2-component Gaussian mixture model fit to the distribution of fluorescence intensity values in a single experiment. Bar graphs show mean ± SEM from n=2 (**C**) or n=3 (**D**, **E**) independent experiments. Scale bars, 50 μm.

### FGF dose sets the proportions of cells with different transcriptional states

In vitro differentiation protocols often yield heterogeneous mixtures of cell types (Jang et al., 2017), and the immunostaining analysis above indicated this was also the case in the mesoderm differentiation protocol applied here. To characterize this heterogeneity, and to determine how it changed depending on FGF concentration, we performed single cell RNA sequencing (scRNAseq) of cells treated with different FGF2 doses from 0 to 12.0 ng/ml (Fig. 3A). In addition, we analyzed cells treated with the selective FGFR inhibitor AZD4547 (FGFRi) during differentiation, to assess possible effects of autocrine and paracrine FGF signaling. We observed massive cell death at micromolar FGFRi concentrations that were previously used in the embryo (Morgani et al., 2018b), and significant cell survival was only obtained at FGFRi concentrations ≤ 31.3 nM (Fig. S1A, B). This prompted us to use a concentration of 30 nM AZD4547 in the scRNAseq experiment. Even though this FGFRi concentration only led to a mild decrease of ERK phosphorylation, a major target of FGF signaling (Fig. S1C), it strongly changed the transcriptional state of the cells as we show below. After quality filtering, we retained approx. 2000 single cell transcriptomes per condition (Table S1). A correlation matrix of pseudo-bulk transcriptomes showed that the samples with a similar FGF signaling strength had the most correlated transcriptomes (Fig. 3B). When comparing neighboring concentrations in concentration series, the biggest pseudo-bulk transcriptomic difference was observed between 0 ng/ml and 0.75 ng/ml FGF2, such that FGF-treated samples clustered away from the non-treated and the FGFRi samples. Analysis of the marker genes T/Bra, Tbx6, Hand1 and Cdx2 confirmed that their concentration-dependent expression that we had detected on the protein level above was recapitulated at the transcriptional level in the scRNAseq dataset (Fig. 3C). We also analyzed the expression of Cdh1 (E-Cadherin) and Cdh2 (N-Cadherin) because cells switch from Cdh1-to Cdh2-expression as they undergo an epithelial-to-mesenchymal (EMT) transition during gastrulation. Cdh1- and Cdh2-expressing cells were found in all conditions, but the fraction of the former decreased, while the fraction of the latter increased with exogenous FGF2. Thus, FGF tunes the proportion of cells undergoing EMT in a dose-dependent manner.

**Fig. 3:**
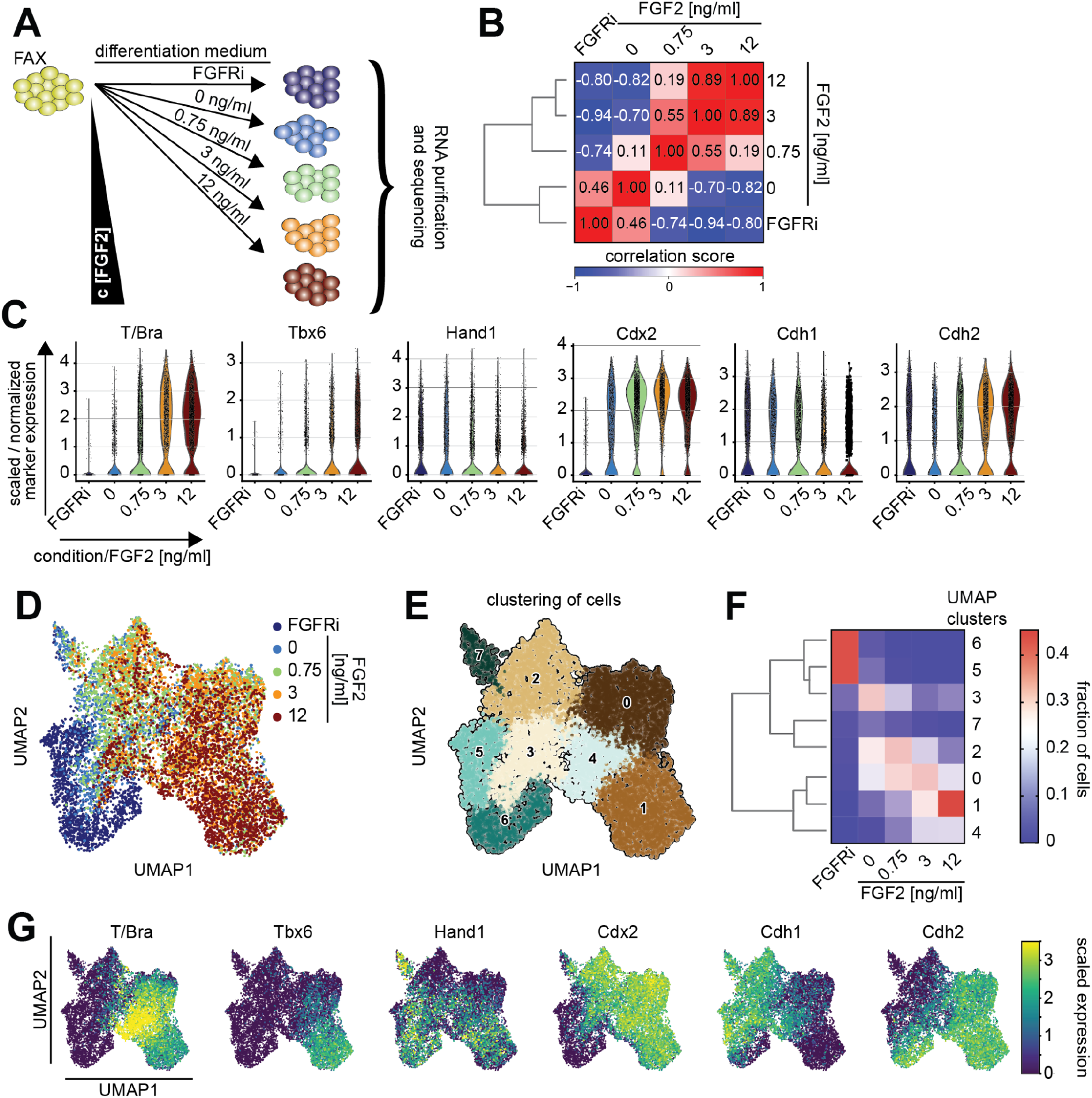
Single-cell transcriptomic analysis of concentration-dependent FGF functions. **A** Schematic of the experimental protocol. EpiSCs from FAX medium were differentiated in the presence of an FGFR inhibitor, without FGF2, or with 0.75 ng/ml, 3 ng/ml, and 12 ng/ml for three days before generation of single-cell transcriptomes and sequencing. The concentrations of Chir and BMP4 were kept at 1 µM and 8 ng/ml as in previous experiments. **B** Correlation matrix of pseudo-bulk samples. Dendrogram on the left shows hierarchical clustering of the samples. **C** Violin plots showing expression levels of marker genes T/Bra, Tbx6, Hand1, Cdx2, Cdh1 and Cdh2 for each of the individual samples. The width of the violin plots is scaled per number of observations. **D** UMAP representation of single cell transcriptomes, color-coded according to treatment regime. **E** Leiden clustering of single-cell transcriptomes from the entire dataset, shown on the UMAP plot from **D**. **F** Heatmap showing the proportion of cells from each sample associated with the clusters identified in **G**. Clusters have been hierarchically ordered based on transcriptional similarity, as indicated by the dendrogram on the left. **G** Expression levels of marker genes T/Bra, Tbx6, Hand1, Cdx2, Cdh1 and Cdh2 color-coded on the UMAP plot from **D**.

To characterize in more detail how the composition of cell types in the population changed with FGF signaling levels, we visualized entire dataset as a dimensionality-reduced Uniform Manifold Approximation and Projection (UMAP) plot (McInnes et al., 2020), and identified transcriptionally similar cells by Leiden clustering (Fig. 3D – F, Table S2). FGFRi-treated cells showed little overlap with cells from the other conditions in the UMAP plot, and were mostly found in clusters 5 and 6. This clear separation of the FGFRi condition from all other samples, including the one without addition of exogenous FGF2, indicates that auto-and paracrine FGF signaling affects gene transcription and cell differentiation. Cells from the other four conditions each occupied different parts of the UMAP plot, and preferentially populated one or two of the 8 clusters (Fig. 3D - F, Fig. S2). However, for each condition, a fraction of cells mapped outside the condition’s main clusters. Since each of the clusters is characterized by the expression of a specific set of marker genes (Fig. 3G, Fig. S3), these observations suggest that, rather than inducing discrete cell states, FGF2 dose shapes the proportions of cells adopting specific states during mesoderm differentiation.

### High exogenous FGF doses promote less advanced cell types

In order to identify the developmental stages and identities of cells differentiated in vitro, and to determine how these changed with exogenous FGF dose, we performed asymmetric dataset integration with *ingest* in Scanpy (Wolf et al., 2018). We used a fully annotated scRNAseq dataset that profiled cells from whole mouse embryos between E6.5 to E8.5 collected at 6 h intervals (Pijuan-Sala et al., 2019), and transferred annotations of developmental stage and cell identity. Projection onto a UMAP representation of the embryo reference data suggested that the heterogeneous cell types differentiated in vitro corresponded to a wide range of cell types and stages present in the embryo (Fig. 4A). Furthermore, cells treated with different doses of FGF2 mapped to different parts of the UMAP plot, indicating that FGF levels affect the representation of cell types and stages (Fig. 4B).

**Fig. 4:**
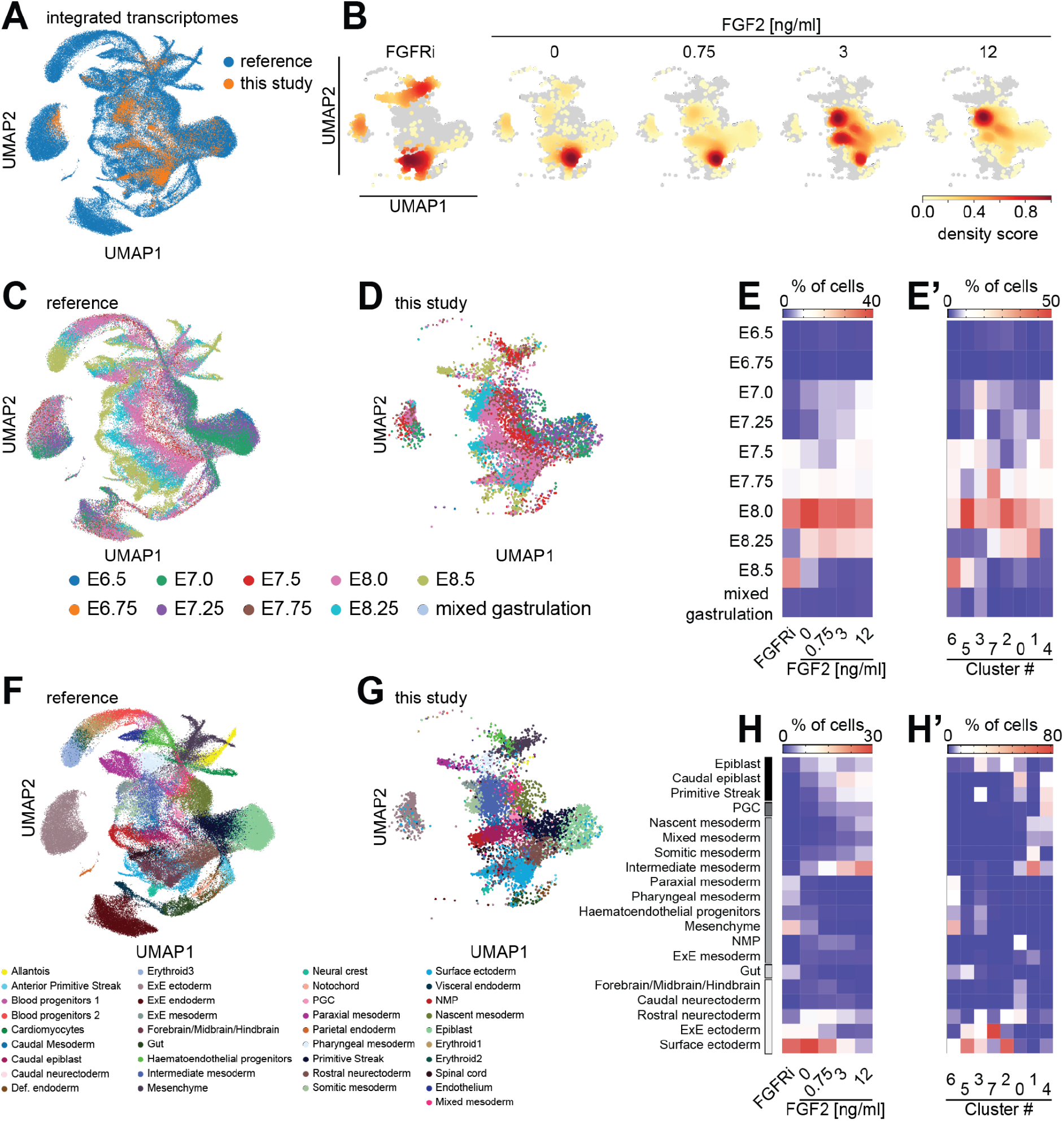
Integration of single cell transcriptomes with an embryo reference reveals developmental stages and cell types obtained in vitro. A UMAP representation of single cell transcriptomes from the reference dataset by Pijuan-Sala et al., 2019 (blue), and the in vitro dataset (orange) after asymmetric integration (see Methods for details). B Transcriptomes from each of the FGF treatment regimes shown separately as density plots, compared to transcriptomes of all other in vitro differentiated cells shown in gray. C - E’ Label transfer to identify developmental stages represented in vitro. C UMAP plot of reference dataset with color-coding according to developmental stage of cells. D UMAP representation of in vitro differentiated cells after integration, color-coded according to stage label transferred from the reference dataset. E Heatmap showing proportion of cells that were assigned a specific stage label for different FGF signaling strengths. E’ Same as E, but showing proportion of cells from each of the clusters identified in Fig. 3E that were assigned a specific stage label. F - H’ Label transfer to identify cell types represented in vitro. F UMAP plot of reference dataset with color-coding according to cell identity. G Same display of transcriptomes from in vitro differentiated cells as in D, but color-coded according to cell-type label transferred from the reference dataset. H Heatmap showing proportion of cells that were assigned a specific cell type label for different FGF signaling strengths. H’ Same as E’, but showing proportion of cells from each of the clusters identified in Fig. 3E that were assigned a specific stage label. Cell types in E, E’, H, H’ are ordered from top to bottom: pluripotent epiblast-like cell types, primordial germ cells (PGCs), mesoderm subtypes, endoderm, and ectoderm-related cell types.

We first analyzed annotations of developmental stages transferred from the embryo reference dataset (Pijuan-Sala et al., 2019, Fig. 4C - E’). Depicting stage labels in the UMAP plot suggested that the in vitro differentiated cells represent a range of developmental stages (Fig. 4D). However, when we plotted the proportion of cells with a specific stage label at different FGF doses as a heatmap we found that for all FGF stimulation regimes, the most prominent stage label corresponded to the E8.0 embryo, followed in most of the conditions by the immediately neighboring stages E7.75 and E8.25 (Fig. 4E). Given that we started from a cell population that is similar to the preimplantation epiblast around E5.0, and then differentiated cells for three days, this observation indicates that differentiation in vitro occurs at a similar pace as in the embryo. However, we also noticed that the FGFRi condition stood out with a large proportion of cells labeled as corresponding to E8.5, while the condition with highest exogenous FGF contained the largest fraction of cells with the stage labels E7.5, E7.25, and E7.0. This shift of stage labels along the FGF axis suggests that high FGF signaling levels can delay differentiation. Consistently, when we analyzed how stage labels were distributed in the clusters determined in Fig. 3E and F, we noticed that clusters 6 and 5 had an overrepresentation of cells with more advanced stage labels, while clusters 3 and 4 were enriched for cells that were classified as less advanced (Fig. 4E’).

To corroborate these findings, we repeated the integration with a second, independent embryo dataset that covered a similar developmental range at lower time resolution (Grosswendt et al., 2020, Fig. S4). Again, the in vitro differentiated cells occupied different areas in the UMAP space when projected on the reference data, in an FGF-dependent manner (Fig. S4A - C). Analysis of the stage labels transferred from this alternative embryo dataset revealed that the large majority of in vitro differentiated cells were classified as corresponding to E7.5 and E8.0 cells, further supporting a similar differentiation pace in vitro and in the embryo. Finally, also here we found a shift of the proportion of stage labels from more advanced stages in the FGFRi condition, to earlier stages in conditions that were treated with high FGF doses (Fig. S4E - F). Taken together, these analyses indicate that heterogeneous cell types emerge during in vitro differentiation with a pace that matches differentiation pace in the embryo, and that exogenous FGF can promote the maintenance of less advanced cell types.

### Dataset integration identifies FGF-dependent and FGF-independent cell types

Next, to better understand which cell types differentiated *in vitro* at different FGF signaling strengths, we analyzed the cell type information obtained by dataset integration. A UMAP representation of the cell type labels transferred from the embryo reference dataset by Pijuan-Sala et al., 2019 suggested that a broad range of cell types were obtained in vitro (Fig. 4G, H). We produced heatmaps to visualize the proportion of cells that were assigned a specific cell type label depending on FGF signaling strength (Fig. 4H), or as a function of the cluster assignments from Fig. 3E and F (Fig. 4H’). We only plotted cell types that contained at least 4% of cells in at least one signaling condition or cluster. To facilitate interpretation of the results, we ordered cell types in the heat maps, starting with epiblast-related cell types at the top, followed by mesoderm, gut, and ectoderm-related cell types. This analysis revealed that epiblast-related cell types were absent in the FGFRi condition, and appeared at low-to-intermediate FGF concentrations (Fig. 4H), suggesting that the maintenance of these pluripotent and developmentally less advanced cell types is promoted by exogenous FGF signals. Inspection of the mesodermal subtypes in the heatmap revealed two broad groups: The first contained cells labeled as “nascent”, “mixed”, “somitic” and “intermediate mesoderm”. These cell types were absent in the FGFRi condition, and their proportion increased with FGF2 dose. The second group contained cells labeled as “paraxial” and “pharyngeal mesoderm”, “hematoendothelial progenitors” and “mesenchyme”, which were found in the FGFRi condition and to a smaller degree in the condition without exogenous FGF, but were nearly absent at FGF2 concentrations ≥ 0.75 ng/ml (Fig. 4H). This suggests that there exist FGF-dependent and FGF-independent mesodermal subtypes. Finally, we observed that ectodermal cell types, in particular cells labeled as “surface ectoderm”, were most prominently obtained in the FGFRi condition as well as in conditions with low exogenous FGF signaling. This observation suggests that a threshold level of FGF signaling is required for converting the previously reported function of BMP4 to induce surface ectoderm differentiation (Li et al., 2013; Malaguti et al., 2013) towards the induction of other lineages. When we plotted the relationship between cell type labels and clusters identified in Fig. 3E, we found that some labels were strongly enriched in specific clusters, such as “intermediate mesoderm” in cluster 1, or “surface ectoderm” in clusters 2 and 5 (Fig. 4H’). In most cases however, clusters contained several related cell type labels, indicating that cluster boundaries in the in vitro dataset do not align with cell type boundaries in the reference dataset.

To substantiate the associations of specific cell types with FGF signaling levels, we analyzed cell type labels that were transferred from the independent embryo dataset by Grosswendt et al., 2020 (Fig. S4G - I’). Here, only few cells labeled as epiblast were detected, in contrast to integration with the dataset from Pijuan-Sala et al., 2019. The proportions of cells with labels corresponding to the primitive streak, different types of mesoderm and ectoderm were broadly consistent between integration with the two datasets. Most notably, we again found two groups of mesodermal labels: one group containing the labels “secondary heart field/splanchnic lateral plate”, “amnion mesoderm early”, “pharyngeal arch mesoderm”, and “hematopoietic/endothelial progenitor” was populated in the FGFRi and the 0 ng/ml condition, whereas the second group containing the labels “primitive streak late”, “presomitic mesoderm”, “NMPs early” and “posterior lateral plate mesoderm” was absent in the FGFRi conditions and increased with FGF2 dose. We notice that the FGF-independent mesoderm cell types identified in vitro are the ones that differentiate first in proximal parts, whereas the mesodermal cell types that we identify as FGF-dependent emerge later and in more distal parts of the gastrula. Thus, this systematic mapping of FGF-independent and - dependent cell types suggests a potential role for FGF signaling in the spatio-temporal patterning of the mesoderm.

### Differentiating cell populations generate divergent endogenous signaling patterns

Having determined how FGF signaling levels affect the proportions of cell types, we next sought to use our data to explore why heterogeneous cell types differentiated within each condition. To test whether this heterogeneity was caused by differential FGF signaling between cell types, we analyzed the expression of the FGF target genes Spry4, Dusp4, Dusp6, and Etv4 (Kalkan et al., 2019; Morgani et al., 2018b, Fig. 5A - C). Expression of these genes was variable across all cells (Fig. 5A), and, as expected, increased with exogenous FGF2 dose (Fig. 5B). FGF target gene expression also differed strongly between clusters (Fig. 5C). This variability of target gene expression between clusters also held true when analyzed separately for each of the FGF stimulation regimes (Fig. 5D). Thus, even upon homogeneous exogenous FGF stimulation, cells show heterogeneous signaling responses. This was not due to differential receptor expression, as the main FGF receptor Fgfr1 was expressed more evenly than the FGF target genes, both across treatment regimes and clusters, and other Fgfrs were not strongly expressed (Fig. 5B - D). However, when we extended our analysis to the expression of endogenous FGFs, we found that expression of Fgf3, Fgf4, Fgf8, and Fgf17 clearly increased with exogenous FGF dose (Fig. 5B). Furthermore, their expression was mainly associated with clusters 0, 1, and 4, both when analyzing the complete dataset (Fig. 5C), as well as within each stimulation regime (Fig. 5D). This indicates that exogenous FGF2 stimulation triggers endogenous FGF signaling. It also raises the possibility that a positive feedback based on short range endogenous FGF signaling amplifies initial heterogeneities locally and eventually leads to the observed cell type heterogeneity.

**Fig. 5:**
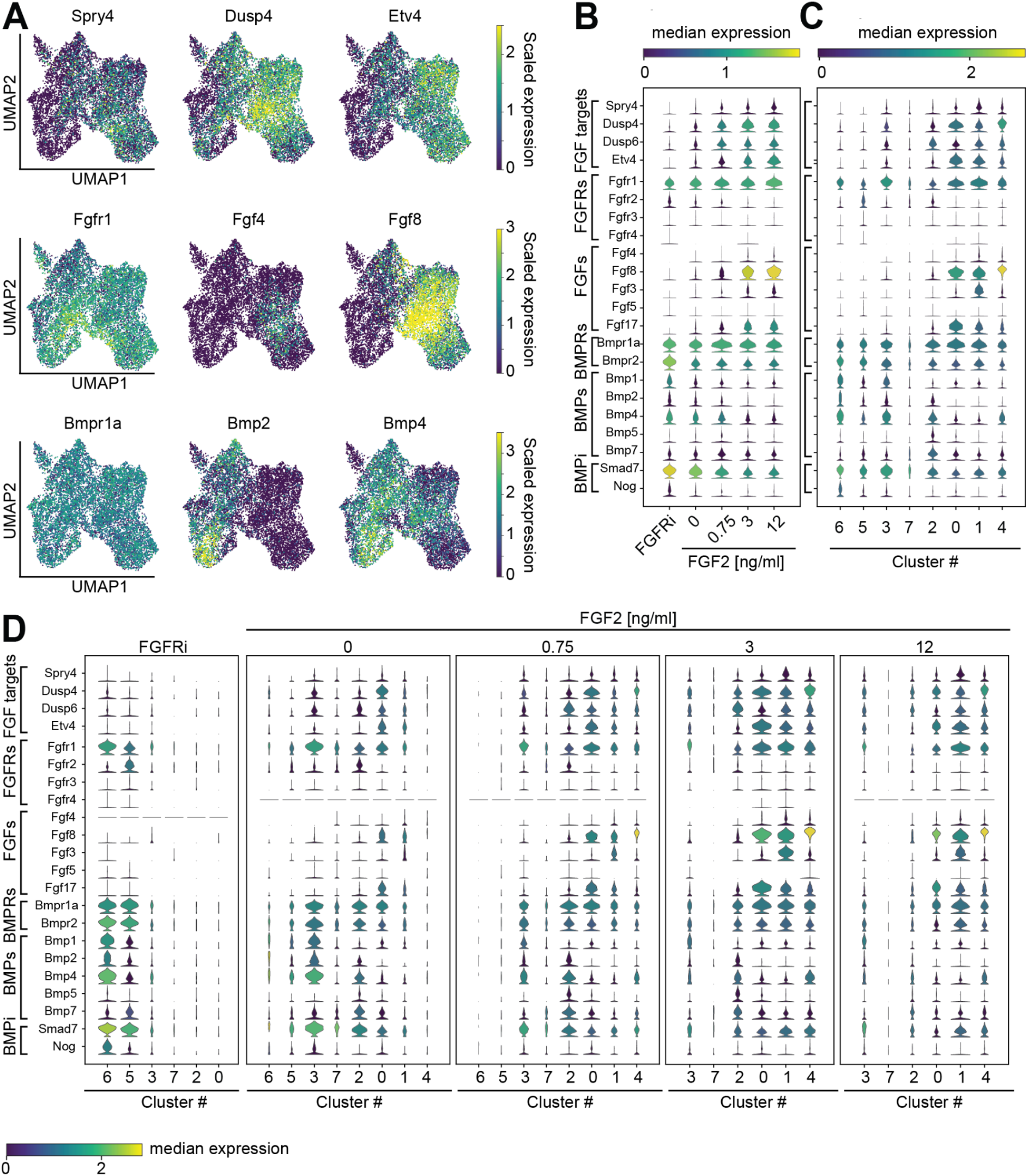
Cluster-specific expression of signaling genes suggests establishment of discrete signaling states in vitro. **A** Single-cell expression of Fgf target genes Spry4, Dusp4, Etv4 (top row), Fgf signaling genes Fgfr1, Fgf4 and Fgf8 (middle row), and Bmp signaling genes Bmpr1a, Bmp2, and Bmp4 (bottom row). Single cells are represented on the same UMAP plot as in Fig. 3C, expression levels of the individual genes are color-coded. **B** Stacked violin plots showing the expression of a panel of genes associated with Fgf and Bmp signaling for each of the FGF signaling regimes. **C** Expression levels of the same genes as in **B**, but segregated according to clusters identified in Fig. 3G. **D** Expression levels of the same genes as in **B**, but divided by treatment condition and grouped by clusters identified in Fig. 3G. Width of the violin plots in **B** - **D** indicates the number of observations, their color reflects the median expression of the selected gene for the cells contained in each observation.

Next, we asked how exogenous FGF affected the expression of BMP, Wnt and Nodal signaling genes. Similar to the pattern of Fgfr1 expression, Bmpr1a was expressed to a similar degree in all experimental conditions and clusters (Fig. 5A - C). The expression of several Bmp ligand genes such as Bmp1, Bmp2, and Bmp4 in contrast decreased with increasing exogenous FGF, and was strongest in clusters 6, 3, and 2 (Fig. 5A - C). Bmpr2 expression also decreased with FGF signaling levels, but its expression was not as cluster-specific as that of the Bmp ligand genes (Fig. 5B, C). The expression of core BMP signaling genes is thus anticorrelated with that of Fgf ligands. This mutually exclusive expression of BMP and FGF ligands by individual cells could act to locally separate their opposing activities in promoting more proximal and more distal fates, respectively.

Finally, we extended our analysis to components of the Wnt and Activin/Nodal signaling systems and found a similar pattern of broad, FGF-independent receptor expression and FGF-regulated, cluster-specific ligand expression (Fig. S5). The Wnt receptor Lrp6 for example was homogeneously expressed in all clusters and conditions, in contrast to several Wnt ligand genes: Wnt3, 4, and 6 expression decreased with exogenous FGF dose, whereas Wnt3a, Wnt5a, and Wnt5b expression increased with FGF2 dose. Expression of Nodal signaling genes was overall low, and expressing cells were predominantly found in clusters 3 and 4. These clusters contain many less differentiated cells (Fig. 4E’, H’), consistent with Nodal’s role in maintaining pluripotency (Guzman-Ayala et al., 2004; Mesnard et al., 2006).

Taken together, our analysis of the expression of signaling genes suggests that populations of differentiating cells can autonomously generate distinct signaling environments.

### Pulsed FGF exposure is sufficient to maintain FGF signaling and T/Bra expression

Finally, we set out to probe the existence of an intercellular positive FGF feedback loop in cell communities, and its role for cell differentiation. We treated cells with FGF2 for one or two consecutive days at different times during the full three-day differentiation protocol (Fig. 6A), and first used the expression of the T/Bra reporter in the SBR line (Deluz et al., 2016) as a read-out FGF-dependent differentiation. Only few reporter-positive cells were detected when cells were exposed to exogenous FGF2 during day 1 or day 3, but FGF treatment during day 2 strongly increased the proportion of reporter-positive cells, to levels similar to those obtained for 2-day and continuous treatment regimes (Fig. 6B, C). This indicates that cells become competent to respond to FGF treatment with upregulation of T/Bra only on day 2. The strong similarity of the proportions of reporter-positive cells in cultures treated for day 2 only compared to cultures treated on both day 2 and day 3 could mean that FGF signaling is only required for a short period of time, or that FGF signaling maintains itself in the population via a positive feedback triggered by pulsed FGF stimulation at day 2. We tested this possibility by pulsed FGF treatment of a Spry4:H2B-Venus reporter cell line that is a faithful transcriptional read-out for FGF signaling (Morgani et al., 2018b). Expression levels of this reporter were similar in EpiSCs growing in the FGF2- containing FAX medium, and in cells grown for 3 days in FGF-containing differentiation medium (Fig. 6D, E). Reporter expression was strongly reduced in differentiating cultures that were not treated with FGF, and those expression levels could not be further reduced by FGFRi treatment (Fig. 6E). Upon pulsed FGF treatment on day 2 (0-F-0 condition), reporter fluorescence reached 69.4 ± 2.9% (mean ± SEM, n=3) for FGF2, and 54.2 ± 7.7% for FGF4 of the value in EpiSCs. However, when the FGF pulse on day 2 was followed by FGFRi treatment on day 3 (0-F-A condition), reporter fluorescence was more than halved compared to the 0-F-0 condition, reaching only 31.8 ± 2.3% for FGF2, and and 22.8 ± 1.9% for FGF4 of EpiSC levels (Fig. 6D, E). This suggests that differentiating cultures maintain FGF signaling following pulsed exposure to FGF. Even though we cannot rule out that residual recombinant FGF bound to the extracellular matrix contributes to the ongoing signaling, this observation is consistent with the activity of a positive intercellular FGF feedback loop in stem cell cultures.

**Fig. 6:**
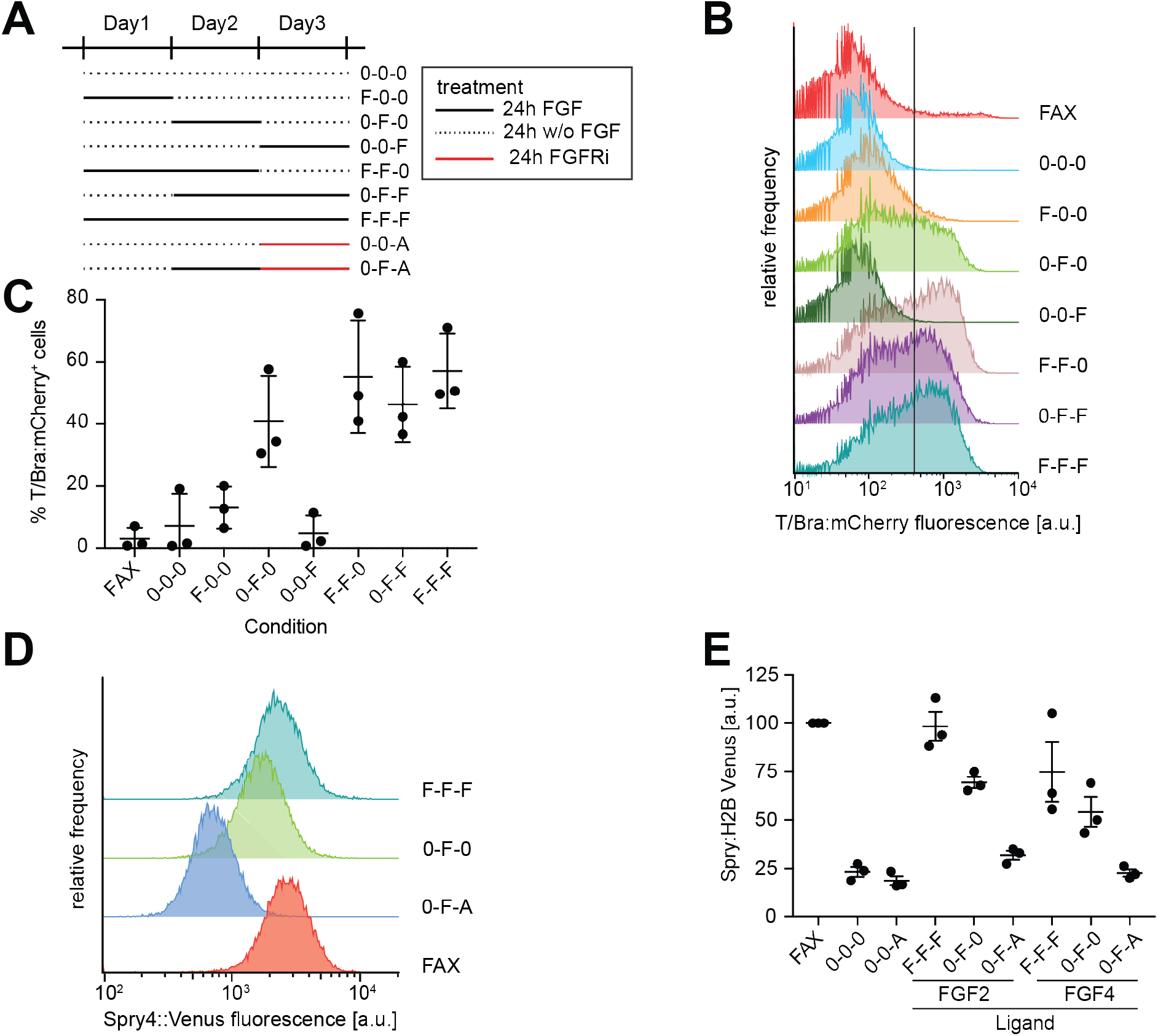
Differentiation and signal transduction upon pulsed FGF treatment. **A** Schematic of timed FGF signaling experiments. Solid black lines represent FGF treatment (F), dashed black lines indicate periods without FGF treatment (0), solid red lines indicate treatment with 30 nM AZD4547 (FGFRi) (A). **B** Representative flow cytometry measurements of T/Bra:mCherry expression in the SBR reporter line differentiated according to the experimental scheme in **A**, using FGF2 as a ligand. Solid line indicates the threshold between reporter-positive and -negative cells. **C** Quantification of percentage of T/Bra:mCherry-positive cells from n=3 independent experiments measured as in **B**. Error bars indicate SEM. **D** Representative flow cytometry measurements of a Spry4:H2B-Venus reporter line differentiated with timed exposure to FGF2. Treatment regimes are labeled according to the scheme in **A**. **E** Quantification of mean Spry4:H2B-Venus fluorescence intensity in cells treated as in **D,** with 6 ng/ml FGF2 or 12 ng/ml FGF4, measured by flow cytometry. Measurements in individual experiments were normalized to the mean intensity in EpiSCs grown in FAX medium. Data from n=3 independent experiments, error bars indicate SEM.

## Discussion

Here, we apply single-cell analysis methods to dissect how FGF and BMP signaling interact to regulate cell type diversity during mesoderm differentiation. We delineate opposing functions of these two signaling systems in mesoderm differentiation, comprehensively identify FGF-dependent and -independent mesodermal cell types, and demonstrate that exogenous FGF dose governs the proportions of cell types. Based on the expression of signaling genes in our single cell dataset we propose an intercellular positive feedback loop centered on FGF signaling, which opposes BMP ligand expression, forming a signaling network that may contribute to mesoderm differentiation and patterning.

By integrating our data with high resolution single-cell transcriptomic maps of embryo development (Grosswendt et al., 2020; Pijuan-Sala et al., 2019), we could characterize the heterogeneous outcome of in vitro differentiation in a more unbiased way and with finer resolution than standard marker-based approaches. Consistent with a previous study that applied a similar approach to scRNAseq data from human ESCs differentiated on micropatterns (Minn et al., 2020), we find that a wide range of embryonic cell types differentiate in vitro. Working with murine cells, the high time resolution of the reference datasets allows us to exactly pinpoint the developmental stage of the in vitro differentiated cells. This analysis indicates that at intermediate FGF doses, differentiation pace in vitro is the same as in the embryo, in contrast to results from a previous study by Morgani et al. which suggested that cell differentiation in vitro is slower than in the embryo (Morgani et al., 2018a). Different starting cell populations may be the cause for these discrepancies. Furthermore, this analysis revealed that FGF signaling levels can impact differentiation speed in vitro, with high FGF levels favoring the maintenance of less differentiated cells, and low FGF leading to more advanced cell types.

Studying mesoderm cell differentiation with stem cells offers opportunities for long-term signaling manipulation that are not available in the embryo. We exploit these opportunities to identify a cohort of proximal mesodermal subtypes such as amnion, mesenchyme, haemato-endothelial progenitors and pharyngeal mesoderm that can differentiate in the absence of FGF signaling, in contrast to mesodermal subtypes that differentiate more distally such as lateral plate, intermediate and somitic mesoderm. A proximo-distal transition from FGF-dependent to FGF-independent mesoderm has initially been suggested from the phenotypes of Fgfr1- and Fgf8-mutant embryos (Ciruna et al., 1997; Deng et al., 1994; Sun et al., 1999; Yamaguchi et al., 1994), but the complexity of the embryonic environment, as well as the interplay of cellular migration and differentiation defects precluded the exact mapping of this transition.

A key result of the present work is that BMP and FGF signaling have opposing functions on the allocation of mesodermal cell fates. Consistent with this, phenotypes arising from loss of BMP-signaling can be rescued by inhibition of FGF signaling (Miura, 2006). How then is this antagonism between these two signaling systems mediated molecularly? Row et al. (2018) have recently proposed a mechanism that operates within single cells through the modulation of bHLH transcription factor activity (Row et al., 2018). Our data suggests an additional - not necessarily exclusive - mechanism that operates via the mutual repression of ligand expression and thus extends the antagonism to the level of local cell communities. Similar regulatory interactions between BMP and FGF signaling and ligand expression have been found in the mouse tail and the zebrafish gastrula (Anderson et al., 2016; Fürthauer et al., 2004), indicating that this may be a general mechanism.

If BMP signaling promotes the specification of earlier-differentiating proximal cell types, and FGF signaling that of later-differentiating distal cell types, then how can mesoderm patterning switch between these two signaling modes? In the embryo, BMP signaling from the ExE triggers Wnt3 expression, which in turn activates T/Bra expression in the epiblast (Ben-Haim et al., 2006), and T/Bra finally targets the Fgf8 promoter (Evans et al., 2012). Thus, FGF signaling in the embryo appears to be activated through a relay mechanism that could create a window of opportunity in which first those cell types differentiate that are BMP-dependent and FGF-independent, until FGF signaling has built up sufficiently to specify more distal cell types. In our 2D mesoderm differentiation protocol, the proportion of FGF-dependent cell types is relatively low in the absence of exogenous FGF, in contrast to the situation in 3D aggregates where FGF-dependent paraxial and axial cell types differentiate efficiently upon pulsed stimulation of Wnt signaling without addition of exogenous FGFs (Brink et al., 2014; Brink et al., 2020). A possible explanation of this difference is that 3D environments may be more conducive to endogenous cell-cell signaling than 2D systems, although we cannot rule out that the continued presence of BMP4 in our protocol inhibited the differentiation of FGF-dependent distal cell types.

Both single cell expression data and timed stimulation experiments suggest that FGF signaling during mesoderm differentiation operates in an intercellular positive feedback loop via the upregulation of ligand genes. This feedback loop could be mediated indirectly via upregulation of Wnt signaling by FGFs (Ciruna and Rossant, 2001; Kelly et al., 2004), followed by the activation of T/Bra expression downstream of Wnt signaling (Arnold et al., 2000; Yamaguchi et al., 1999), and finally activation of Fgf8 expression by T/BRA (Evans et al., 2012). Fgf ligands and Fgf target genes tend to be expressed in the same restricted groups of cells despite widespread Fgfr1 expression, suggesting that this positive feedback loop operates locally. Spatially restricted, positively autoregulatory cell-cell communication is the basis for the community effect (Bolouri and Davidson, 2010; Saka et al., 2011). The community effect describes a situation where a critical number of cells in close contact produce and respond to a paracrine signal, ultimately leading this group of cells to differentiate along the same lineage (Gurdon, 1988; Gurdon et al., 1993a). The differentiation of muscle progenitors in Xenopus, a cell lineage related to the somitic mesoderm that differentiates at high FGF2 concentrations in our work, has been proposed to rely on a community effect based on eFGF (Gurdon et al., 1993b; Standley et al., 2001). We therefore speculate that an analogous FGF-based community effect may orchestrate mesoderm differentiation in mammals.

Populations of pluripotent stem cells have a remarkable potential to self-organize into reproducible patterns of differentiated cell types, both in 2D and 3D (Fu et al., 2021; Liu and Warmflash, 2021; Warmflash et al., 2014). It is clear that these processes must be coordinated by cell-cell interactions through mechanical and chemical signals. Most studies to explain self-organized patterning in stem cell models for development have focused on the intercellular organization of the Wnt, BMP and Nodal signaling systems (Chhabra et al., 2019; Etoc et al., 2016; Tewary et al., 2017). Our findings on how FGF signaling is regulated within populations, and how it interfaces with other signaling systems calls for including these connections in future models for cell differentiation and patterning during mammalian gastrulation.

## Methods

### Cell culture

EpiSCs were routinely grown in FAX medium on fibronectin-coated tissue culture plastic. Coating was performed with 20 µg/ml human fibronectin (Merck FC010) in PBS for at least 30’. FAX medium is N2B27 supplemented with 12 ng/ml FGF2 (Cell Guidance System GFM12), 25 ng/ml ActivinA (Cell Guidance System GFM029) and 20 µM XAV939 (Cell Guidance System SM38). N2B27 was prepared as a 1:1 mixture of DMEM/F12 (PAN Biotech P04-41150) and Neuropan basal medium (PAN Biotech) with 0.0025% bovine serum albumin (Gibco), 1× N2 and 1× B27 supplements (Thermo Fisher Scientific), 1x GlutaMAX (Gibco), and 50 μM 2-mercaptoethanol. Cells were split with Accutase (Merck) every two days or upon reaching confluency, and replated at a density of 8000 to 10000 cells per cm^2^. For mesoderm differentiation, EpiSCs were seeded at a density of 1300-2000 cells/cm^2^. The day after seeding, medium was changed to N2B27 supplemented with growth factors and small molecule inhibitors: CHIR99201 (Merck), murine BMP4 (PeproTech 315-27), FGF2 (Cell Guidance System), FGF4 (PeproTech), or AZD4547 (Selleckchem). During differentiation, medium was changed daily.

### Cell lines

The wild type EpiSC line used in this study is from a 129 background and was obtained from Dr. Jennifer Nichols. This line has been derived from the epiblast of an E6.5 embryo by culturing on fibronectin-coated dishes in standard EpiSC medium, consisting of 12 ng/ml FGF2 and 25 ng/ml Activin A in N2B27. The SBR Sox1-T/Bra reporter cell line, and the Spry4 H2B-Venus reporter cell line have previously been described (Deluz et al., 2016; Morgani et al., 2018b). Both lines were transitioned from their original culture media to an EpiSC state by culturing for at least 8 passages in FAX medium. All cell lines were regularly tested for mycoplasma contamination.

### Immunostaining

Cells for immunostainings were cultured on ibidi 8-chamber µ-slides (ibidi), and processed as previously described (Schröter et al., 2015). Primary antibodies used were anti-Nanog 1:200 (Thermo Fisher Scientific, eBioMLC-51), anti-T/Bra 1:200 (R&D, AF2085), anti-Hand1 1:200 (R&D, AF3168), anti-Tbx6 1:200 (R&D, AF4744), and anti-Cdx2 1:250 (BioGenex, MU392A-5UC). Secondary antibodies coupled with appropriate AlexaFluor dyes were from ThermoFisher, and used at 1:500 dilution.

### Imaging and image analysis

Live cells were imaged on an Axiovert 40 brightfield microscope equipped with a Leica MC 170 HD camera. Fluorescence imaging was performed on a Leica Sp8 confocal microscope with a 63x oil immersion objective. Bigger fields of view were imaged using the multi-tile settings of the Leica software, with 10% overlap between single tiles, followed by stitching with the BigStitcher plug-in in FIJI (Hörl et al., 2019; Schindelin et al., 2012).

Nuclei were segmented with the FIJI plug-in StarDist (Schmidt et al., 2018), using a custom-trained model. Training data for the model was obtained by first generating masks of Hoechst-stained nuclei with the pre-trained StarDist model ‘*Versatile (fluorescent nuclei)*’, followed by manual correction. The custom model was then trained on the StarDist 2D ZeroCostDL4Mic platform (v 1.12, Chamier et al., 2021) for 40 epochs on 7 paired image patches, starting from the pretrained model ‘*Versatile (fluorescent nuclei)*’. This custom-trained model was used on the Hoechst channel to determine ROIs corresponding to individual nuclei. To check for correct segmentation, the ROIs were projected onto the nuclear channel, and any segmentation mistakes were manually corrected. The ROIs were then used to retrieve fluorescence intensities for each of the marker channels. On average, 8 independent fields of view were analyzed per experiment and condition, to capture the spatial variability of the expression of the chosen differentiation markers. The data were further analyzed and plotted with custom scripts written in Python. Thresholds between marker-expressing and non-expressing cells were determined by fitting a two-component Gaussian mixture model to the distribution of fluorescence intensities. Cells were considered marker-positive if they belonged to the component with the higher expression levels with a greater than 70% probability, otherwise they were considered marker-negative.

### RT-qPCR

Cells for RT-qPCR analysis were detached from the culture vessel using Accutase, centrifuged and either processed immediately or snap-frozen in liquid nitrogen for long-term storage at -80°C. RNA was isolated from cell pellets with TRIZOL (Thermo Fisher Scientific) according to the manufacturer’s instructions. Following spectrophotometric quality control and quantification, the RNA was diluted to a concentration of 100 ng/μl and stored at -80°C. RT-qPCR was performed with the Luna Universal One-Step RT-qPCR Kit (New England Biolabs) according to manufacturer’s instruction on an iQ5 Real-Time PCR System (Bio-Rad). Primer pairs used are given in Table S3. For each primer pair, we determined the efficiency of amplification with a titration series of template. For each sample, we ran three replicates per condition for the housekeeping genes and two replicates for the marker genes. The fold-change in transcript levels was calculated for each primer pair as *efficiency^−(Ct(condition) − Ct(EpiSC))^*. To determine this fold-change, we used the same Ct(EpiSC) for all experiments which had been determined on an independent plate. The fold-change of each marker gene was then normalized by dividing through the geometric mean of the fold-change of the two housekeeping genes. Finally, to compare concentration-dependent behavior between transcripts, we normalized values for each transcript within a concentration series to their highest expression value.

### Western blotting

Cell samples for immunoblotting were washed twice with ice-cold PBS supplemented with 1mM orthovanadate, and then incubated with lysis buffer. Lysis buffer was prepared fresh by supplementing a commercially available cell lysis buffer (Cell Signaling) with complete EDTA-free protease inhibitor cocktail (Roche), phosphatase inhibitor cocktail 2 and 3 (Sigma) and benzonase (Thermo Fisher Scientific). Cell lysis was aided by scraping, and lysates were collected and snap-frozen in liquid nitrogen. Samples were then analyzed for quality control and quantification with a micro-BCA assay (Thermo Fisher Scientific). Lysates were denatured using a standard Laemelli buffer and boiled for 5 minutes at 95°C. Samples were then incubated into ice for 5 minutes, and between 10 µg and 20 μg protein was loaded on a Bis-Tris SDS gel. Gels were run with 1x MOPS buffer (Thermo Fisher Scientific) with fresh sodium bisulphite. After the run, the gels were transferred on methanol-activated PVDF membranes (Millipore) at 40 V for 90 minutes with a NuPage transfer system (Thermo Fisher Scientific) and incubated with primary and secondary antibodies for 1 h each. The membranes were imaged with the Odyssey Infrared Imaging System (LI-COR Biosciences). Primary antibodies used were anti-Tubulin 1:5000 (Sigma, T6074), anti- pERK1/2 1:1000 (Cell Signaling, 4370S), and anti-total ERK1/2 1:1000 (Abcam, ab36991). Secondary antibodies used were IRDye-coupled donkey anti-mouse 800 at 1:500 and donkey anti-rabbit 680, both at 1:500 dilution (LI-COR Biosciences).

### In-Cell Western

To obtain the high cell densities required in In-Cell Western experiments, 7000 cells/cm^2^ were seeded in fibronectin-coated 96-well black plates with transparent polystyrene bottom (Corning). Fixation, blocking, and incubation with primary antibodies were performed as described for immunostaining experiments above. Primary antibodies used were anti-Hand1 1:500 (R&D, AF3168), anti-Tbx6 1:500 (R&D, AF4744), anti-Gata6 1:1000 (R&D, AF1700), and anti-Tubulin 1:5000 (Sigma, T6074). Secondary antibodies IRDye800-conjugated donkey anti-goat and IRDye680- conjugated donkey anti-mouse (LI-COR Biosciences) were used at 1:500 dilution and incubated in the dark for 1 h. Fluorescence signals were acquired with an Odyssey Infrared Imaging System (LI-COR Biosciences), and regions of interest corresponding to individual wells were selected in the images with the Fiji plugin MicroArray Profile. Signal intensities for proteins of interest were normalized against the anti-Tubulin signal.

### Flow cytometry

Cells for flow cytometry were cultured in 6-well plates, detached with Accutase, centrifuged, and either resuspended in PBS supplemented with 1% BSA for immediate analysis, or fixed in 4% PFA at room temperature for 15 minutes, followed by centrifuging and resuspending in PBS with 1% BSA. Before analysis, cells were passed through a 40 µm cell strainer to obtain single cells and remove cell clumps. Reporter expression in the SBR line was measured in live cells on a BD FACSAria Fusion flow cytometer (BD Biosciences). Spry4:H2B-Venus reporter cells were fixed before measurement, and analyzed on a BD LSRII flow cytometer (BD Biosciences). At least 20.000 cells were analyzed per condition. Data was further analyzed in FlowJo (BD Biosciences).

### Single Cell RNA Sequencing

Single cell transcriptomes were generated with the 10x Genomics Chromium Next GEM Single Cell 3ʹ Reagent Kit v3.1 according to the manufacturer’s instructions. Briefly, cells were differentiated for three days, detached with Accutase and counted. The cell suspension was diluted to a concentration of 900 cells/μl, to recover around 2000 cells per sample. *Cell suspensions were loaded on a Chromium Controller (10x Genomics) to partition cells with gel beads in emulsion. Reverse transcription, cDNA recovery and amplification, and sequencing library construction were performed according to manufacturer’s instructions (10x Genomics ChromiumNextGEMSingleCell_v3.1_Rev_D). We chose 12 PCR cycles for cDNA amplification, and 10 PCR cycles for index PCR. Concentration and fragment size of sequencing libraries were determined with a BioAnalyzer High Sensitivity DNA Assay (Agilent). Libraries were sequenced by paired-end Illumina sequencing on a NovaSeq6000 instrument with a read length of 150 bp. We first performed sequencing at shallow depth with a target of* 20*10^6^ reads per sample*, to estimate the number of cells captured in each sample, and to confirm the generation of high-quality single-cell transcriptomes. In a subsequent deeper sequencing run, we adjusted sequencing depth for each sample depending on captured cell number, to* a depth between 125*10^6^ and 180*10^6^ reads per sample*. Analysis was carried out on data from the deep sequencing run only*.

Demultiplexing, alignment to the mouse genome mm10 (GENCODE vM23/Ensembl 98, from 10x Genomics) and read quantification was performed with CellRanger (10x Genomics, v4.0.0). The libraries were saved as annotated dataset objects (anndata) with extension ‘.h5ad’. All further analyses were carried out with the Python package Scanpy 1.7.0rc1, Wolf et al., 2018, and https://scanpy.readthedocs.io/en/stable/tutorials.html). Briefly, we first performed quality control and preprocessing on each of the 5 anndata files retrieved from CellRanger separately, filtering out barcodes that had less than 2500 genes per cell, or more than 10% of unique molecular identifiers (UMIs) mapping to mitochondrial genes. Following concatenation, the resulting dataset was normalized to a value of 50’000 reads per cell and log-transformed. We selected for highly variable genes, and regressed out effects of total counts per cell and the percentage of mitochondrial genes expressed. Finally, the data for each gene was scaled to unit variance, and single count values exceeding standard deviation 10 were clipped. Dimensionality reduction was performed by first running a Principal Component Analysis, followed by neighborhood analysis and visualization in two dimensions using the Uniform Manifold Approximation and Projection technique (McInnes et al., 2020). Leiden clustering (Traag et al., 2019) was performed with a resolution of 0.33. Hierarchical clustering and calculation of pearson correlation scores of aggregated, pseudo-bulk transcriptomes from each condition was performed with the function “scanpy.tl.dendrogram” in Scanpy, using highly variable genes. Scanpy was also used to produce density plots, and to visualize the expression of specific genes as violin plots or in the UMAP plot. To produce heatmaps, tabular data from Scanpy were plotted in Prism 7 (GraphPad).

### Dataset integration

Our own single cell transcriptome data was integrated with two independent published single cell transcriptome datasets covering the development of the mouse embryo from E6.5 to E8.5. Data from Pijuan-Sala et al. (Pijuan-Sala et al., 2019) was retrieved in R with the Bioconductor experimental data package “MouseGastrulationData” available at https://bioconductor.org/packages/release/data/experiment/html/MouseGastrulationData.html. Data from Grosswendt et al. (Grosswendt et al., 2020) was obtained from the authors as an R object. Both datasets were translated into a Python *anndata* object, and processed as described above for our own scRNAseq data. Dataset integration was performed with the *Ingest* function of *Scanpy,* using exclusively genes that were common between the reference embryo dataset and the *in vitro* dataset. We used the integration to transfer labels describing developmental stage and cell type identity from the reference dataset.

## Acknowledgements

We thank Jenny Nichols and David Suter for sharing cell lines, Philippe Bastiaens and Ivan Bedhov for helpful suggestions throughout the project, and Melania Barile and Max Fernkorn for help with scRNAseq analysis. We are grateful to Helene Kretzmer and Stefanie Grosswendt for sharing processed scRNAseq data. We thank members of the Schröter lab, David Turner and Melania Barile for comments on the manuscript, and the IMPRS-CMB for support.

## Data and code availability

The single cell sequencing data from this study have been deposited at GEO with accession number GSE228061. The full documentation and code of the scRNAseq and image analysis is available at https://github.com/Schroeterlab/mesoderm_diff.

---

**Fig. S1:**
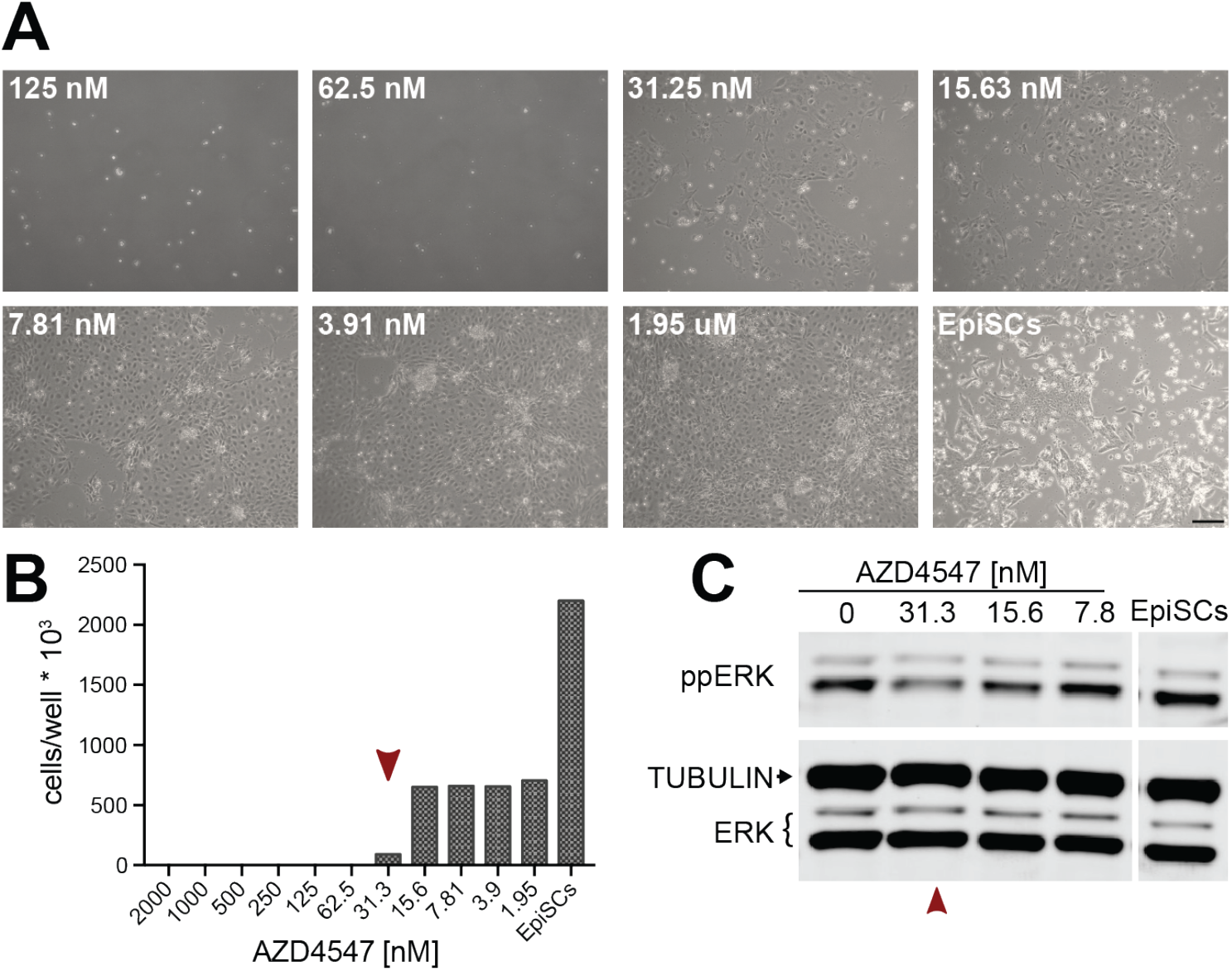
Titration of the FGFR inhibitor AZD4547 during mesoderm differentiation. **A** Brightfield images of cells after differentiation for 3 days with 1 μM Chi, 8 ng/ml BMP4 and indicated concentrations of AZD4547. Scale bar, 250 µm. **B** Cell count after treatment as in **A**. **C** Immunoblotting for ppERK and total ERK of cells treated for 24 h with 1 μM Chiron, 8 ng/ml BMP4 and indicated concentrations of AZD4547.

**Fig. S2:**
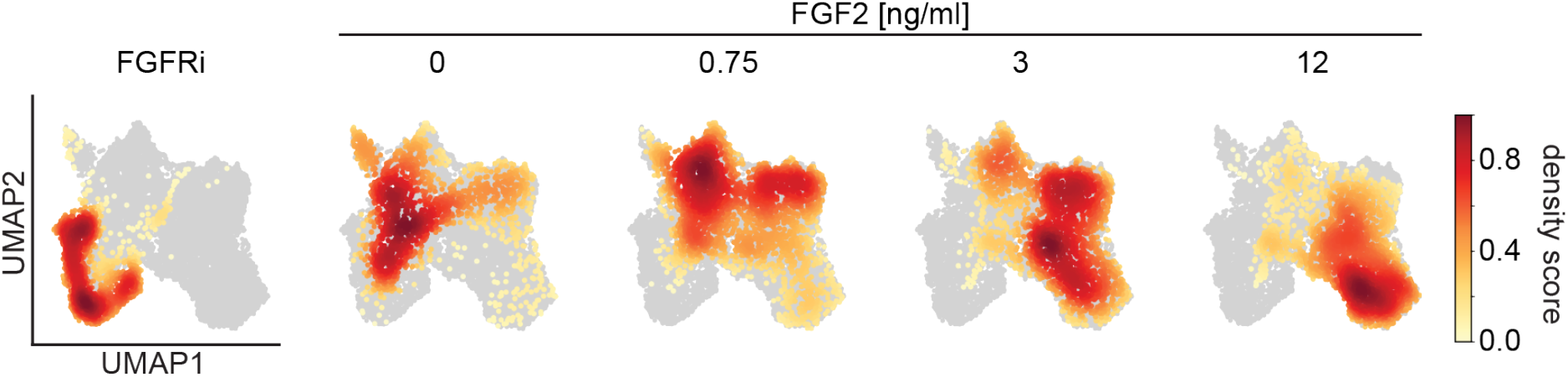
Distribution of cells from individual samples in UMAP space. Density plots showing the distribution of cells from each sample in the UMAP plot from Fig. 3D. Remaining cells from the dataset are shown in gray.

**Fig. S3:**
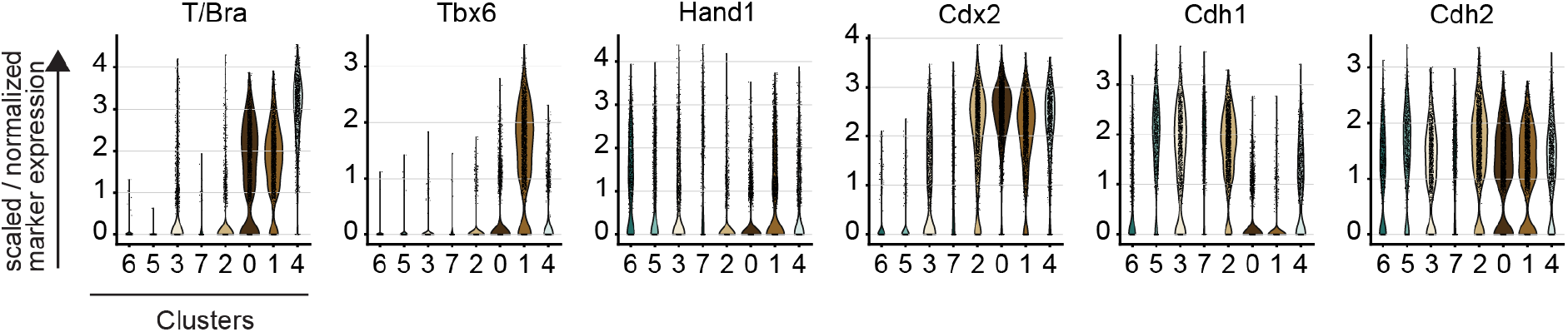
Expression of marker genes in cell clusters identified by scRNAseq. Violin plots showing the expression levels of T/Bra, Tbx6, Hand1, Cdx2, Cdh1 and Cdh2 for clusters identified in Fig. 3G. The width of the violin plots is scaled per number of observations.

**Fig. S4:**
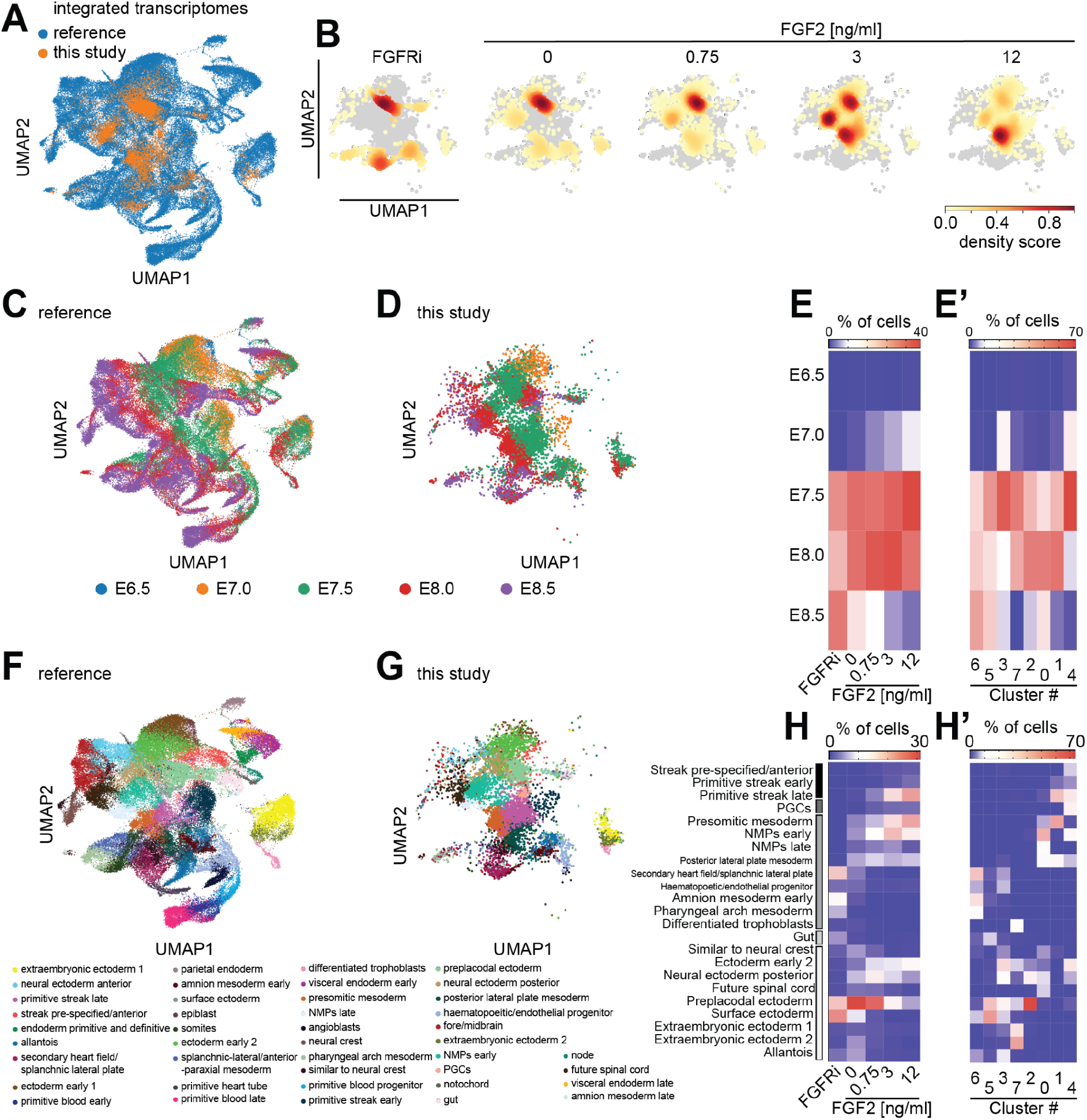
Integration of single cell transcriptomes with an alternative embryo reference reveals developmental stages and cell types obtained in vitro. **A** UMAP representation of single cell transcriptomes from the reference dataset by Grosswendt et al., 2020 (bue) and the in vitro dataset (orange) after asymmetric integration. **B** Transcriptomes from each of the FGF treatment regimes shown separately as density plots, compared to transcriptomes of all other in vitro differentiated cells shown in gray. **C - E’** Label transfer to identify developmental stages represented in vitro. **C** UMAP plot of reference dataset with color-coding according to developmental stage of cells. **D** UMAP representation of in vitro differentiated cells after integration, color-coded according to stage label transferred from the reference dataset. **E** Heatmap showing proportion of cells that were assigned a specific stage label for different FGF signaling strengths. **E’** Same as **E**, but showing proportion of cells from each of the clusters identified in Fig. 3G that were assigned a specific stage label. **F - H’** Label transfer to identify cell types represented in vitro. **F** UMAP plot of reference dataset with color-coding according to cell identity. **G** Same display of transcriptomes from in vitro differentiated cells as in **D**, but color-coded according to cell-type label transferred from the reference dataset. **H** Heatmap showing proportion of cells that were assigned a specific cell type label for different FGF signaling strengths. **H’** Same as **E’**, but showing proportion of cells from each of the clusters identified in Fig. 3G that were assigned a specific stage label. Cell types in **E**, **E’**, **H**, **H’** are ordered from top to bottom: pluripotent epiblast-like cell types, primordial germ cells (PGCs), mesoderm subtypes, endoderm, and ectoderm-related cell types.

**Fig. S5:**
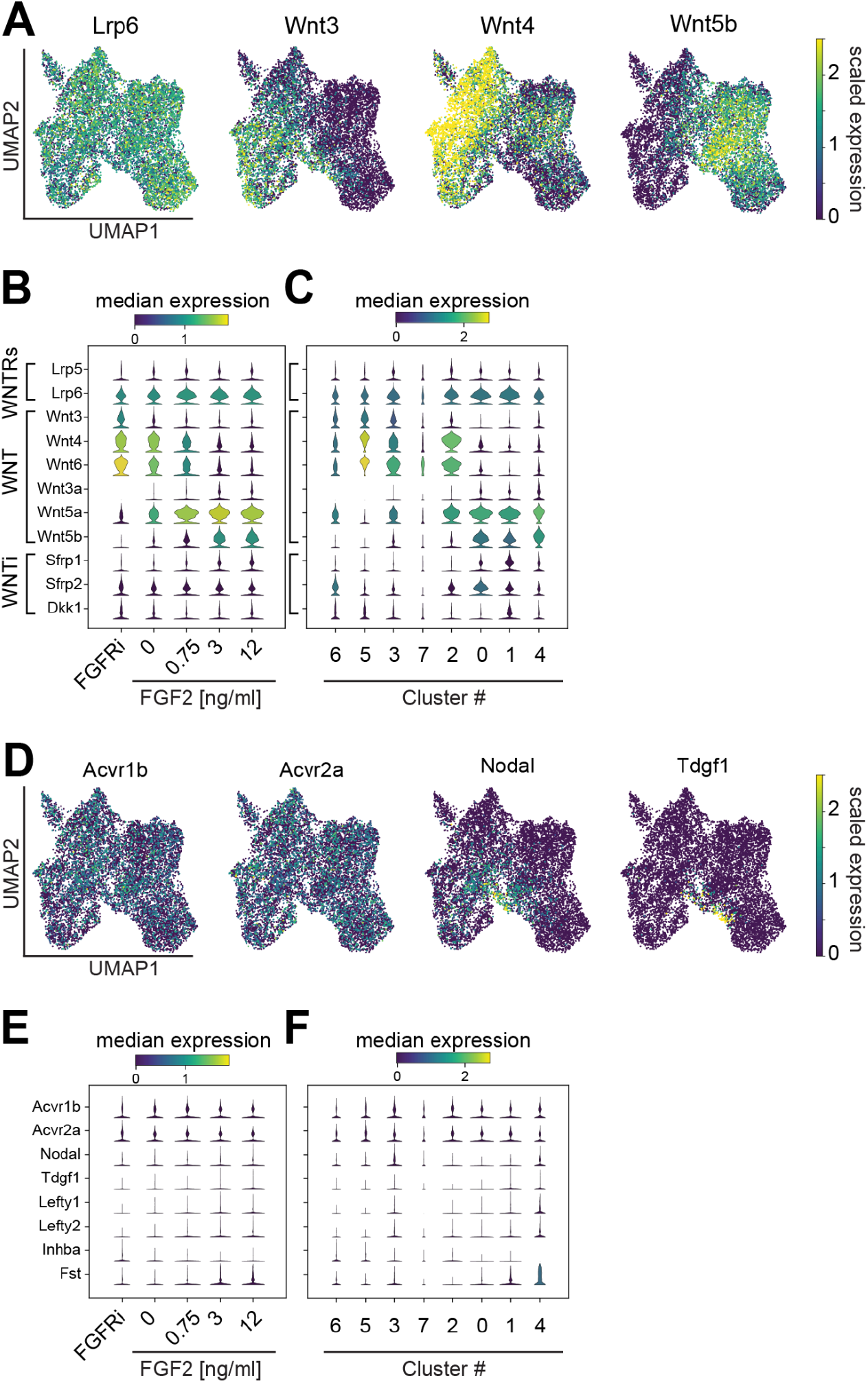
FGF- and cluster-specific expression of Wnt and Nodal signaling genes. **A** Single-cell expression of the Wnt receptor Lrp6 and ligands Wnt3, Wnt4, and Wnt5b. Single cells are represented on the same UMAP plot as in Fig. 3C, expression levels of the individual genes are color-coded. **B** Stacked violin plots showing the expression of a panel of genes associated with Wnt signaling for each of the FGF signaling regimes. **C** Expression levels of same genes as in **B**, but segregated according to clusters identified in Fig. 3G. **D** Same as **A**, but for the Nodal receptors Acvr1b and Acvr2a, and the ligands Nodal and Tdgf1. **E**, **F** Same as **B**, **C**, but for a panel of genes associated with Activin/Nodal signaling. Width of the violin plots in **B**, **C**, **E**, **F** indicates the number of observations, their color reflects the median expression of the selected gene for the cells contained in each observation.

**Table S1.** Summary statistics of single cell sequencing dataset.

| Sample | number of cells | average number of genes | average number of reads | % reads mapped to mitochondrial genes |
| --- | --- | --- | --- | --- |
| all | 10653 | 5294 | 30185 | 4.3 |
| FGFRi | 1701 | 5088 | 28116 | 4.5 |
| 0 ng/ml FGF2 | 1696 | 5472 | 33498 | 4.3 |
| 0.75 ng/ml FGF2 | 2725 | 5277 | 29923 | 4.1 |
| 3 ng/ml FGF2 | 2125 | 5362 | 30403 | 4.3 |
| 12 ng/ml FGF2 | 2406 | 5273 | 29417 | 4.3 |

**Table S2.** Table shows number of cells from every experimental condition that was allocated to each of the eight clusters identified in Fig. 3E.

|  | Cluster |  |  |  |  |  |  |  |
| --- | --- | --- | --- | --- | --- | --- | --- | --- |
| Sample | 0 | 1 | 2 | 3 | 4 | 5 | 6 | 7 |
| FGFRi | 17 | 0 | 23 | 111 | 1 | 757 | 758 | 34 |
| 0 ng/ml FGF2 | 334 | 110 | 435 | 515 | 14 | 116 | 52 | 120 |
| 0.75 ng/ml FGF2 | 778 | 320 | 851 | 429 | 228 | 18 | 3 | 98 |
| 3 ng/ml FGF2 | 661 | 557 | 359 | 146 | 378 | 0 | 0 | 24 |
| 12 ng/ml FGF2 | 453 | 1089 | 201 | 218 | 435 | 0 | 0 | 10 |

**Table S3.**
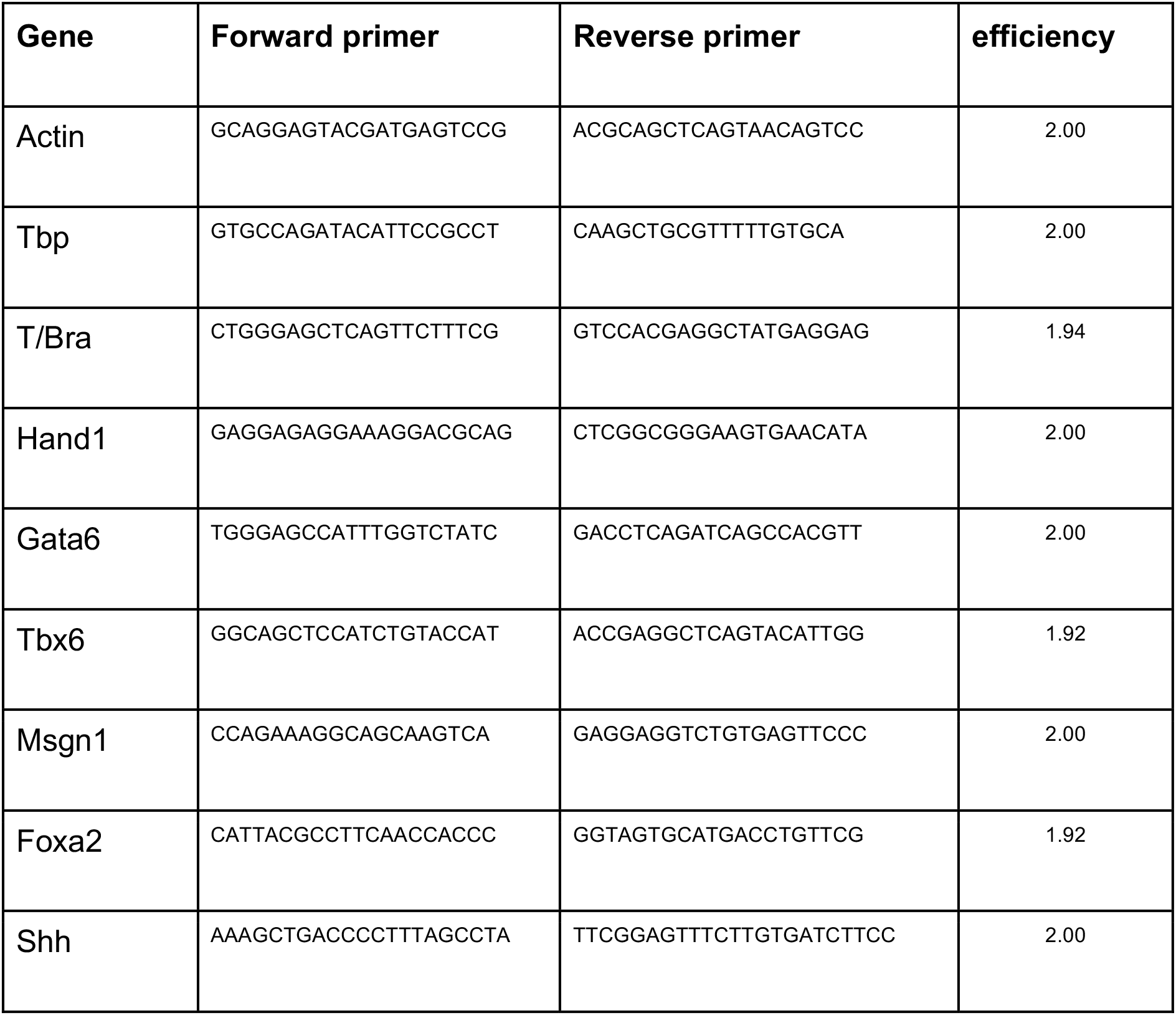
Sequences of primers used in RT-qPCR experiments.

